# High Cellular Uptake Gene Delivery Platform with Chitosan and L-arginine Complex for Cancer Treatment

**DOI:** 10.1101/2022.02.22.481395

**Authors:** Maidou Wang, Hongqian Zhang, Chuanxu Yang

## Abstract

Gene therapy outperforms chemotherapy in limiting the harm to human body. A major impediment to the progress of gene therapy is a lack of proper carriers. Chitosan, derived from shellfish, is a desirable candidate for nucleic acid delivery because of its outstanding biodegradability and biocompatibility. However, previous works show that chitosan has poor cell membrane permeability. In this study, we synthesize a complex delivery system for gene therapy by compounding chitosan with L-arginine, and test the encapsulation, uptake, and cancer cell treatment power. Encapsulation efficiency of the chitosan/L-arginine complex (CA) can reach more than 90% in weight. Flow cytometry reveal that siRNA delivered by the complex can be taken in by cells 10 times better than free siRNA and 2 times better than chitosan alone. Confocal imaging confirmed the high cellular uptake, and siRNA tests in HeLa cells indicate the successful silencing of the target gene (RRM2). The complex serves as a gene delivery system that is capable of loading and delivering a variety of genetic materials in which the specific nucleic acid molecule to be delivered could be altered at will.

## 1. Introduction

Globally, cancer is the second leading cause of death, killing millions of people every year.[1] Conventional chemotherapy is most often prescribed as a treatment of metastatic cancer, but it still has many unwanted side-effects, arising from low selectivity and drug resistance.[2] Chemotherapy functions by inhibiting certain kinds of enzymes or proteins to disrupt the fast-growing mechanism inside cells. Thus, chemotherapeutic agents kill not just the cancer cells, but all cells that rapidly divide. This could cause severe damages to many of the essential but fast-growing tissues, such as blood, digestive tract, or bone marrow.[3] Besides, cancer cells obtain resistance against chemotherapy over time. Through DNA mutations and metabolic adjustments, cells gradually develop multiple drug-resisting mechanisms, such as drug inactivation, DNA damage repair, and cell death inhibition etc., significantly reducing the drug’s anticancer effect and leading to chemotherapy failure.[4]

Gene therapy, on the other hand, has the potential to get around all obstacles that chemotherapy faces. Gene therapy acts by using genetic molecular substances to either repair or impair the genetic process inside cells [5]. There are several mechanisms by which gene therapy works, including replacing a dysfunctional gene with its healthy counterpart, silencing a disease-causing gene that produces too much of an unwanted substance, and inserting a new gene that possesses particular functions, such as triggering apoptosis or enhancing immune cell recognition. [6] From these basic mechanisms, specific branches of gene therapy are derived, such as antisense oligonucleotides (ASOs), which deactivates or eliminates certain non-functioning mRNAs by delivering its complementary strand [7], RNA interfering, which inhibits certain overly-expressed genes by preventing their corresponding mRNAs from undergoing translation [8], and suicide gene therapy, which introduces suicide genes to code for proteins or enzymes that induce cell death [9]. Gene therapy is superior to ordinary chemotherapy methods because it circumvents the problem of low specificity and remarkably lessens on drug resistance. Gene therapy has high specificity since it exclusively targets at the disease-causing gene, strictly distinguishing cancer cells from ordinary fast-dividing cells.[10] It even indirectly achieves specificity by transducing drug-resistant genes into normal cells, protecting healthy tissues against systematic chemotherapy. [11] Drug resistance can be lowered in gene therapy. With its gene-inactivation mechanisms, gene therapy could knock down the resistance gene to restore sensitivity.[12] [13]

In spite of all the potential advantages, gene therapy development is yet in an onset stage. Still more needs to be done to achieve large-scale production and avoid side-effects like immune responses, long-term storage instability, and insertional mutagenesis.[14] Many of the existing downsides of gene therapy come with the choice of gene carrier. For example, about 70% of gene therapy clinical trails use viral vector for delivery,[15] while it is almost inevitable that a viral vector would evoke immune responses.[16]

With the hope of finding a more superior carrier, we conducted the following study.

RRM2 was chosen as our target. RRM2 stands for Ribonucleotide reductase regulatory subunit M2. It encodes ribonucleotide reductase that serves as a catalyst during the production of genetic materials. This gene is often selected as the typical prognostic target in quite a number of cancers such as liver cancer, [18] breast cancer, [19] and prostate cancer[20]. Overexpression of RRM2 also interact with the signaling pathways that control the cell chemoresistance level of cancer. [21] Thus, inhibition of RRM2 gene can effectively enhance the anticancer effect of chemotherapeutic drugs and elicit cancer cell apoptosis, further controlling the growth of tumor.

RNA interference is a promising technology in which small interfering RNA (siRNA) carries out targeted mRNA cleavage and, subsequently, suppresses the expression of specific genes.[22] Hence, we will adopt siRNA as therapeutic agent to silence RRM2 and lead to death of tumor cells.

However, there are several barriers to natural siRNA’s clinical applications, such as low membrane permeability,[23] and sensitivity to nuclease etc.[24]. Meanwhile, considering that the carrier must also avoid incurring immune responses, high membrane permeability and biocompatibility become two key criteria of a suitable carrier.

Chitosan (Cs) and its derivatives, obtained from the exoskeleton of shellfish, are desirable nucleic acid carriers. Chitosan is completely harmless to human body. Coming from a natural source of Chitin, chitosan can be degraded by human body and thus considered biodegradable, and its degraded molecular materials do not interfere with normal human body functions and thus considered biocompatible. Reports have also confirmed its low toxicity and does not arouse overreaction of human immune system. Chitosan holds positive charge because of its amine groups and can easily compound with genetic materials through electrostatic interactions because DNA and siRNA holds negative charges[25].

Nevertheless, Cs by itself is not a perfect candidate for siRNA delivery, due to the reported limited cell membrane-permeability[26]. Here in this work, we added L-arginine onto the chitosan framework to form a complex for enhanced membrane penetration, and at the same time retaining the strong tendency to bind with genetic molecules and maintain its biosafety. Figure 1 schematically illustrated the design idea. Because of its strong positive charge[27], arginine has been reported to show an exceptional membrane permeability.[28] Arginine-rich Cell Penetrating Peptides also have predominantly better membrane permeability.[29] Thus, arginine was attached to the Cs carrier to further improve the extent of intake by cells. The Cs/L-arginine compound holds stronger positive charges than pure Cs and therefore it tends to interact with siRNA stronger and form compound nanoparticles more easily. Meanwhile, since arginine is one of the essential amino acids that human body needs, the complex would be safe for human body.

**Figure 1.**
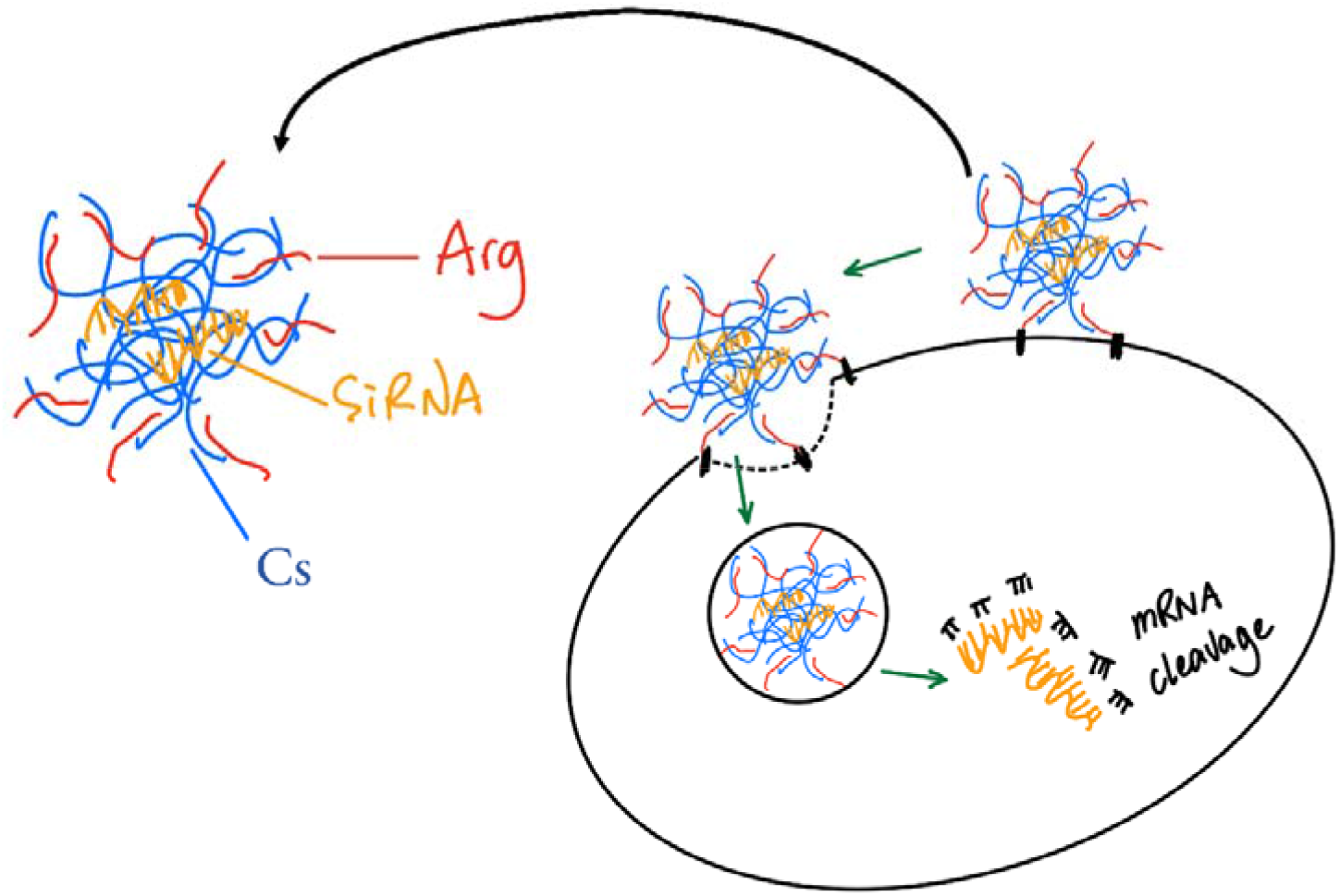
Schematic drawing of the formation of Cs/L-arginine complex encapsulating siRNA for gene delivery

To summarize, Cs was modified with the addition of arginine, obtaining a Cs-Arg compound that acts as siRNA carrier. The carrier possesses lower immunogenicity compared with nonviral vectors, and better membrane permeability than natural Cs.

## 2. Materials and Methods

### 2.1 Synthesis of chitosan (Cs) and chitosan/L-arginine(CA) complex

The chitosan(Cs)/L-arginine complex in our work was synthesized by mixing Cs powder and L-arginine in the EDC/HCL-HOBT buffer with a mass ratio 2.5 (Cs):1.0 (L-arginine). The pH of the buffer was adjusted to neural or acidic (=<7.0) with HCl. Nanoparticle carrier of Cs was synthesized in HOBT buffer by mixing 250mg Cs with 13mL buffer. HOBT buffer was synthesized by dissolving 250mg of Hydroxybenzotriazole (HOBT) into 13mL of DEPC water. EDC/HCL-HOBT buffer was prepared by mixing 250mL of HOBT buffer with 400mg of EDC. (EDC=1-(3-Dimethylaminopropyl)-3-ethylcarbodiimide hydrochloride)

### 2.2 Preparation of Delivery System

To synthesize the delivery system siRNA-Cs/CA, we designed and ordered 21bp siRNA sequence and loaded the siRNA into Cs/CA nanoparticles with self-assembly reactions. The 21bp-double-stranded siRNA molecules were assembled by putting together two complementary strands of 21bp-single-stranded siRNA molecules. The resulting siRNA molecules were diluted to 20 μM/L using DEPC water.

The delivery system with chitosan (Cs) nanoparticles and the chitosan-arginine (CA) nanoparticles were assembled by combining 21bp-double-stranded siRNA molecules and Cs/CA complexes at various weight ratios. Cs powder or dried CA complexes were then dissolved in HOAc-NAOAc buffer at a concentration of 1mg/mL. Dissolved Cs/CA was blended with diluted siRNA at weight ratios of 1, 5 and 10 (Cs/CA to siRNA) to react. All reactions took place in the HOAc-NaOAc buffer. HOAc-NaOAc buffer was prepared using HOAc of 200mM/L and NaOAc of 200mM/L with a volume ratio of 14 to 86 (HOAc: NaOAc). Buffer was finalized to pH=5.45 (±0.05) (adjusted with HCl or NaOH). The weight ratio was adjusted by fixing the siRNA concentration (20μM/L) and varying the Cs/CA concentration. The mixture of Cs/CA complexes and siRNA was placed in room temperature for 30 minutes to fully react.

### 2.3 Encapsulation efficiency

The siRNA-Cs/CA mixture after reaction was added to TE buffer, which was synthesized using Tris-HCl of 1 M/L, EDTA of 0.5M/L and H2O with a volume ratio of 5: 1: 400 (Tris-HCl: EDTA: H2O), in accordance with the manufacturer’s instructions. The pH of TE buffer was adjusted to 5.4. The encapsulation efficiency was quantified with fluorescence. siRNA was marked with Ribogreen fluorescence marker, which was diluted in TE buffer with 1:1000 ratio (excitation: 500 nm; emission: 525 nm). Untreated siRNA free in solution buffer was included as the blank compare group. Every sample measurement repeated 3 times in 3 different hole-wells on a 96-well plate, and the final encapsulation efficiency was the average of the three repeating measurement. The encapsulation efficiency was calculated by averaging out the values of the same sample in 3 distinct wells. Tecan Spark microplate reader was utilized to measure fluorescence.

### 2.4 Preparation for cellular uptake measurement

For cell culture, we prepared approximately 50,000 cells in each well (24-well plate), and then incubated at human body temperature (37) for a day (24 h). siRNA had been marked with Cy5 for flow cytometric identification. Dissolved Cs/CA was blended with diluted siRNA at weight ratios of 5 and 10 (Cs/CA to siRNA) in the HOAc-NAOAc buffer and the complete medium. The weight ratio was adjusted by fixing the siRNA concentration at 50mM/L and varying the Cs/CA concentration. The samples were put into the wells, with two repetitions for each sample. The mixtures of cells and samples were kept in 37 degree Celsius for 4 hours. Cultural medium was removed. Cells were washed with PBS. Then we added Trypsin to dissolve them. The mixture of cells and trypsin was kept in 37 degree Celsius for 1 minute to dissolve and then PBS buffer containing 2% FBS was added. Finally Cells were remixed and re-dispersed. Cells were then remixed and pipetted to spread out in the solution. The mixtures were then taken to flow cytometry.

### 2.5 Confocal Imaging

Cells were directly nurtured in a 4-chamber confocal plate at 37 degree Celsius for 24 hours, in which 50,000 cells were put into each well. Samples of free siRNA, Cs-siRNA nanoparticles (Cs : siRNA weight =10:1), and CA-siRNA nanoparticles (CA : siRNA weight =10:1) were distributed into the wellplate. The mixtures of cells and samples were kept in 37 degree Celsius for 4 hours, after which the old culture medium was replaced with new culture medium, and the blended system were kept in 37C for 1h. All liquid was extracted, leaving the cells behind. The cells were washed three times using PBS. 500 uL of 4% paraformaldehyde was put into each chamber, after which the blended system was kept in 37C for 15 minutes. All liquid was extracted, leaving the cells behind, which were washed three times using PBS. 500uL of Hoechst (2ug/mL) was put into each chamber, then the blended system was kept in 37C for 10 minutes. All liquid was extracted, leaving the cells behind, which were washed three times using PBS. Anti-fluorescence quenching agent was added to the cells. A Leica TCS SP8 was used for taking confocal images of the cell samples.

### 2.6 FTIR and NMR preparations and measurements

To measure FTIR, we used a Bruker Tensor II model installed in Shandong University facility center. To prepare samples for FTIR measurement, first prepare 100mg dry KBr powder in the container, add 2mg sample and grind the mixture under infrared light gently until powder looks finely grounded and mixed well. Load the mixture into pressure and compress the mixture to obtain nearly transparent thin flask.

To obtain NMR measurement, prepare 8mg of sample and dissolve completely in 1 mL buffer. The buffer is made from deuterated water and deuterated acetic acid with volume ratio 99 to 1. The 1H-NMR was measured at 400 MHz on a NMR spectrometer model AVANCE 400 from Bruker.

### 2.7 Apparatus

Fluorescence intensity was measured on a Tecan Spark microplate reader. Flow cytometry was conducted on a flow cytometer of NovoCyte model from ACEA Biosciences. Inc. To measure FTIR, we used a Bruker Tensor II model. All machines and apparatus are installed in Shandong University facility center.

### 2.8 Materials

Chitosan powder (95%, deacetylation from chitin) and L-arginine was purchased from Aladdine Industrial Corporation (Shanghai, China). Cy5 and Hoechst dye were purchased from Sigma Aldrich. Chemicals used for buffer preparation, such as Hydroxybenzotriazole, 1- (3-Dimethylaminopropyl)-3-ethylcarbodiimide hydrochloride, HOAc, NaOAc were acquired from Aladdine Industrial Corporation (Shanghai, China). All chemicals were analytical grade pure (>97%). siRNA was ordered and purchased from GenePharma company. Cell line used for siRRM2 was HeLa cells, purchased from Procell company.

## 3. Results

### 3.1 Synthesis / Characterization of Cs and CA complex

Cs/L-arginine(CA) complex can be used as a drug carrier for cell intake. The complex in our work was synthesized by mixing Cs and L-arginine in EDC/HCL-HOBT buffer with a mass ratio 2.5 (Cs):1.0 (L-arginine). For control experiments, nanoparticle carrier purely composed of Cs was also synthesized in HOBT buffer. After the reactions, Cs powder and dried Cs/L-arginine powder were taken to FTIR measurement after FTIR preparation. In the FTIR range of wavenumber (min=500cm-1, max=4000cm-1), FTIR measurements can show absorptions features related to specific molecular structures due to the molecular vibrations and thus it can be used to tell if L-arginine was successfully linked onto Cs nanoparticles.

For pure Cs, in the spectrum from 3750 cm-1 and 2500cm-1, we identified a wide hump band indicated active R-OH hydroxyl groups jutting out from the monomer molecules of chitosan polymer[30]. In the range of 1750 cm-1 to 1500 cm-1, there are two notable peaks, a and b, as denoted on the graph. Peak a corresponds to special double bonds between carbon atoms in the amide group, which is the leftover from deacetylation of chitin to make chitosan. Peak b indicates the presence of amino groups, which is unique to pure chitosan and seldom found in chitosan-arginine as chitosan connects with arginine by losing one hydrogen atom from this amino group. With the attachment of L-arginine, the CA complex shows feature similar to that of the Cs complex in the wavelength range of 3750 cm-1 to 2500cm-1, while it also displays different features in the wavelength range of 1750 cm-1 to 1500 cm-1. Peak **c** indicates the presence of guanido group, which exists on arginine but not on chitosan. Peak **d** indicates the presence of amide bond, which is formed through condensation reaction between the hydroxyl group of arginine and the amino group of chitosan. The spectrum proved that L-arginine is indeed attached to chitosan, and that the chitosan-arginine complex is successfully formed.

1H-NMR spectrum was taken to reveal the chemical environment of the hydrogen atoms in the samples. The feature of the hydrogen atoms can also be used to tell if the L-arginine was successfully connected to chitosan. Several peaks are seen in both Cs and CA in the range between 3.9 ppm and 2.9 ppm, denoted on Figure 2 by **a, b, c, a’, b’ and c’**. They represent the hydrogen atoms on the chitosan backbone, confirming the presence of chitosan in the CA complex. Peak **d** is exclusively seen in CA in the range between 2.8 ppm and 2.7 ppm. It represents the methylene groups in arginine, confirming the successful attachment of L-arginine to chitosan.

**Figure 2.**
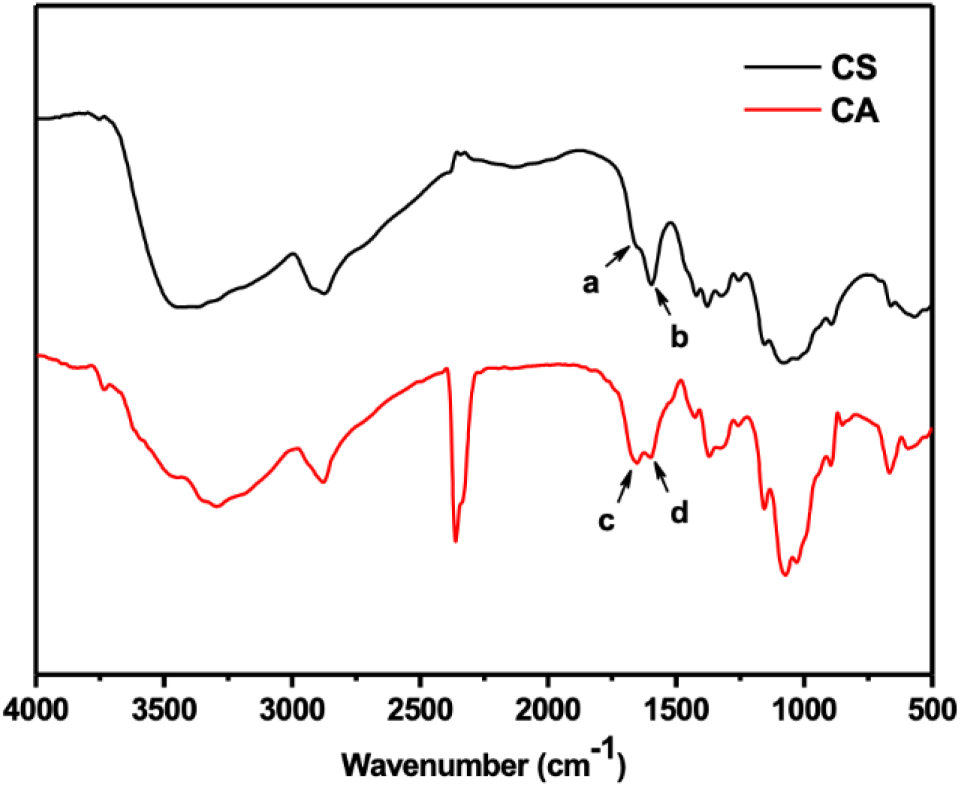
FT-IR (Fourier Transform InfraRed) spectrum of Cs (Chitosan complex, black curve) and CA (Chitosan complex with L-Arginine attached, red curve)

### 3.2 Encapsulation Efficiency

#### 3.2.1 Preparation of Nanoparticles

The delivery system comprised of a specially designed double stranded siRNA and chitosan (Cs) nanoparticles and the chitosan-arginine (CA) nanoparticles. The nanoparticles were assembled by combining the 21bp-double-stranded siRNA molecules and Cs/CA complexes at various weight ratios. Cs powder or dried CA complexes were then dissolved in HOAc-NAOAc buffer and blended with diluted siRNA at weight ratios of 1, 5 and 10 (Cs/CA to siRNA) to react. The weight ratio was adjusted by fixing the siRNA concentration (20μM/L) and varying the Cs/CA concentration. The siRNA-Cs/CA mixture was placed in room temperature for 30 minutes to fully react before measurement of encapsulation efficiency.

#### 3.2.2 Encapsulation efficiency

siRNAs were marked with fluorescence markers to help evaluate the efficiency of encapsulation by nanoparticle carriers. When the siRNA strands are free in solution, they can fully absorb and re-emit light. However if they are fully “encapsulated” into the Cs or CA complex, the energy of the absorbed light would transfer from the genetic molecule to the Cs or CA nanoparticle compound and therefore no re-emitting light would be measured. This so-called “quenching” effect can be used to determine how much siRNA was captured by the nanoparticle carriers by measuring the downsize of light intensity re-emitted by siRNA-Cs/CA. siRNA was marked with Ribogreen fluorescence marker, which was diluted in TE buffer with 1:1000 ratio (excitation: 500 nm; emission: 525 nm). We used untreated siRNA free in solution buffer as the blank compare group.. Every sample measurement repeated three times in three different hole-wells on a 96-well plate, and the final encapsulation efficiency number was the average of the three-repeating measurement.

In Figure 3, the encapsulation efficiency for both Cs and CA complexes are over 90% at all weight ratios and not significantly changed with higher proportions of Cs or CA in the reactions. For CA, the efficiency of weight ratio 1:10 is the highest at 96%, while the efficiency of weight ratio 1:1 is the lowest at 92%. The results suggested that our CA compound has high capacity in encapsulating siRNA for drug delivery purpose. The high encapsulating capacity is probably the result of having high amount of positive charge from both chitosan and L-arginine.

**Figure 3.**
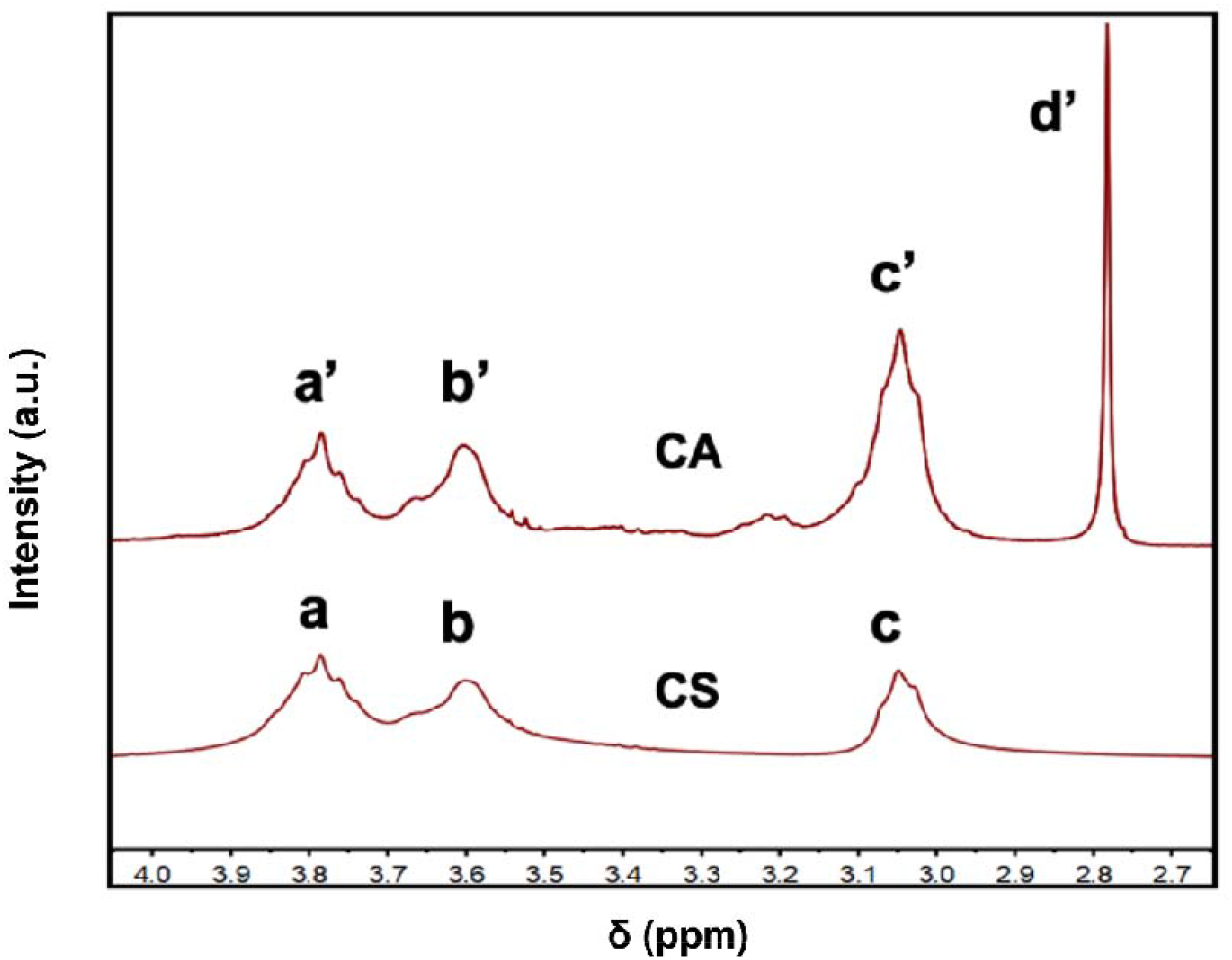
1H-NMR spectrum of Cs (Chitosan complex, bottom curve) and CA (Chitosan complex with L-Arginine attached, top curve)

**Figure 4.**
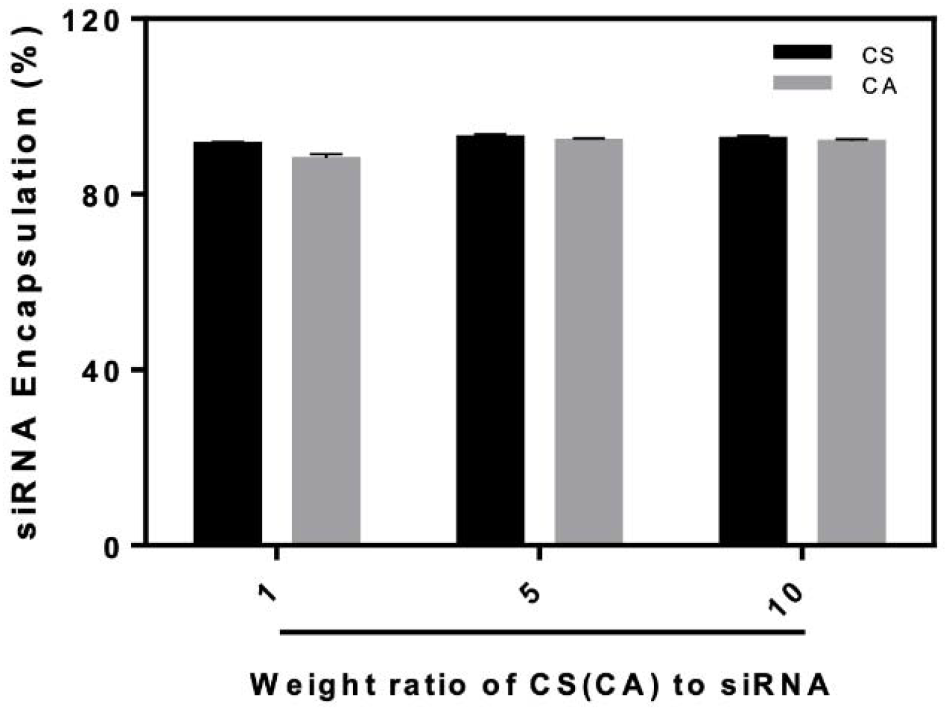
Encapsulation efficiency of Cs (Chitosan complex) and CA (Chitosan complex with L-Arginine attached) for siRNA at different weight ratios of Cs(CA) to siRNA.

**Figure 4.**
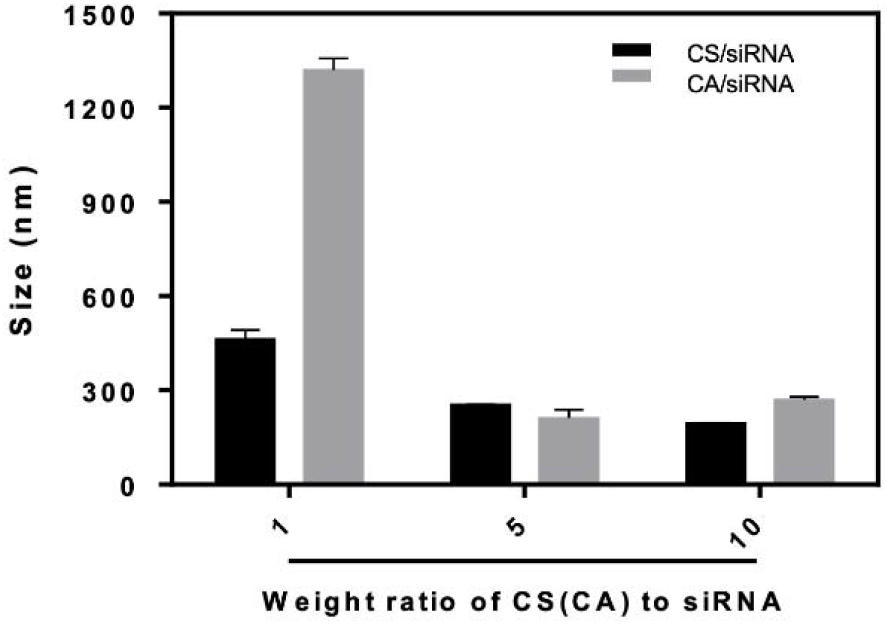
Size of Cs (Chitosan complex) and CA (Chitosan complex with L-Arginine attached) for siRNA at different weight ratios of Cs(CA) to siRNA.

### 3.3 Hydrodynamic diameter

The siRNA-Cs/CA mixture after reaction was added into TE buffer for this measurement. The particle size of the siRNA-Cs/CA nanoparticles was measured using Zetasizer Nano analyzer. Particles with three different Cs/CA-to-siRNA weight ratios had been analyzed. The sizes of different nanoparticle variations are obtained by averaging out the sizes of all nanoparticles with the same variation.

The nanoparticle diameters are bigger at lower weight ratio (Cs/CA : siRNA = 1:1) and smaller at higher weight ratios (5:1 and 10:1). At 1:1 weight ratio, the CA particle diameter is over 1200 nm, too large for cellular uptake. At weight ratio 5:1 or 10:1, the diameters of both the Cs nanoparticles and the CA nanoparticles are below 300 nm, suitable for cellular intake. At these two weight ratios, the Cs and CA plus DNA nanoparticle sizes do not have a big difference. Specifically, when it comes to the CA nanoparticles, the diameter is comparatively smaller when the Cs/CA-to-siRNA weight ratio is 5:1, as opposed to 10:1. Given that particle diameters below 300 nm are preferred for cell uptake, we use weight ratio 5:1 and 10:1 for the next experiment test.

### 3.4 Cellular Uptake with Flow cytometry

Cells were put into a 24-well plate with approximately 50,000 cells in each well and kept in 37 degree Celsius for 24 hours. siRNA had been marked with Cy5 for flow cytometric identification. Dissolved Cs/CA was blended with diluted siRNA at weight ratios of 5 and 10 (Cs/CA to siRNA) in the HOAc-NAOAc buffer and the complete medium. The weight ratio was adjusted by fixing the siRNA concentration at 50mM/L and varying the Cs/CA concentration. The mixtures were carefully re-dispersed before taken to flow cytometry. The Figure 5(A) shows the intensity of Cy5 within live cells, reflecting the success of the cellular uptake of siRNA. For untreated cells in which no siRNA is present, the mode fluorescence intensity is 10^3, and the average fluorescence intensity is almost negligible. For cells exposed to free siRNA without Cs/CA carriers, the mode fluorescence intensity is 10^5, and the average fluorescence intensity is still almost negligible. For cells exposed to Cs nanoparticles with a Cs-to-siRNA weight ratio of both 5 and 10, the mode fluorescence intensity is 5.0 * 10^5, and the average fluorescence intensity also 5.0 * 10^5. For cells exposed to CA nanoparticles with a CA-to-siRNA weight ratio of 5, the mode fluorescence intensity is 9.0 * 10^5, and the average fluorescence is also 9.0 * 10^5. For cells exposed to CA nanoparticles with a CA-to-siRNA weight ratio of 10, the mode fluorescence intensity is 2.0 * 10^6, and the average fluorescence is 2.3 * 10^6. Looking at both the mean and the mode fluorescence intensity, cellular uptake is better for Cs nanoparticles and CA nanoparticles in both weight ratios, compared with free siRNA, indicating that the Cs/CA carrier has played a decisive role in facilitating cellular uptake of siRNA. Comparing between Cs and CA nanoparticles, the mean fluorescence intensities for CA carriers are higher than that for the Cs carriers. It is obvious that the fluorescence intensity is stronger for CA nanoparticles in both weight ratios than Cs nanoparticles, indicating that the addition of L-arginine can evidently improve the nanoparticles’ membrane permeability, as expected. Cellular uptake is significantly better with a higher CA complex weight ratio in the mix, proving that the nanoparticle is better delivered into cells when the CA-to-siRNA weight ratio is closer to 10.

**Figure 5.**
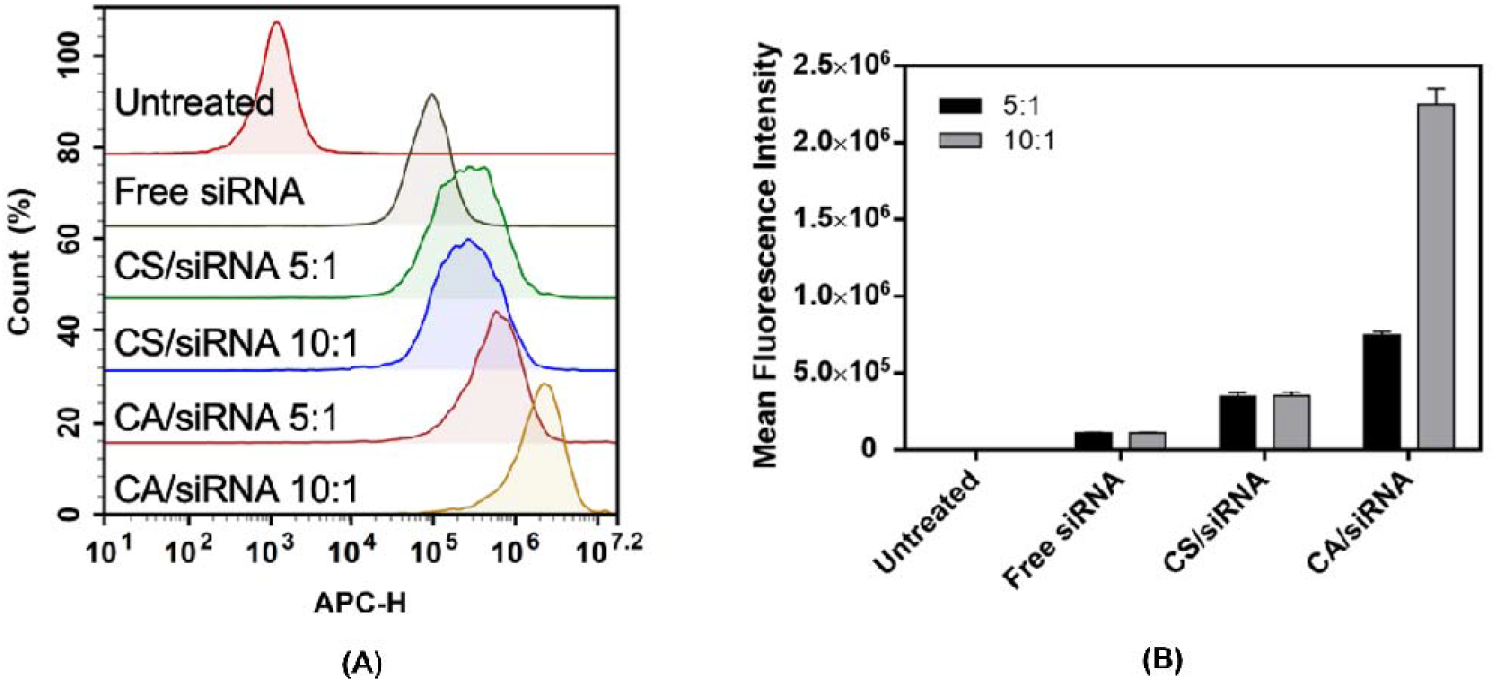
Flow cytometry analysis of Cs nanoparticles and CA nanoparticles at different weight ratios of Cs(CA) to siRNA.

**Figure 6.**
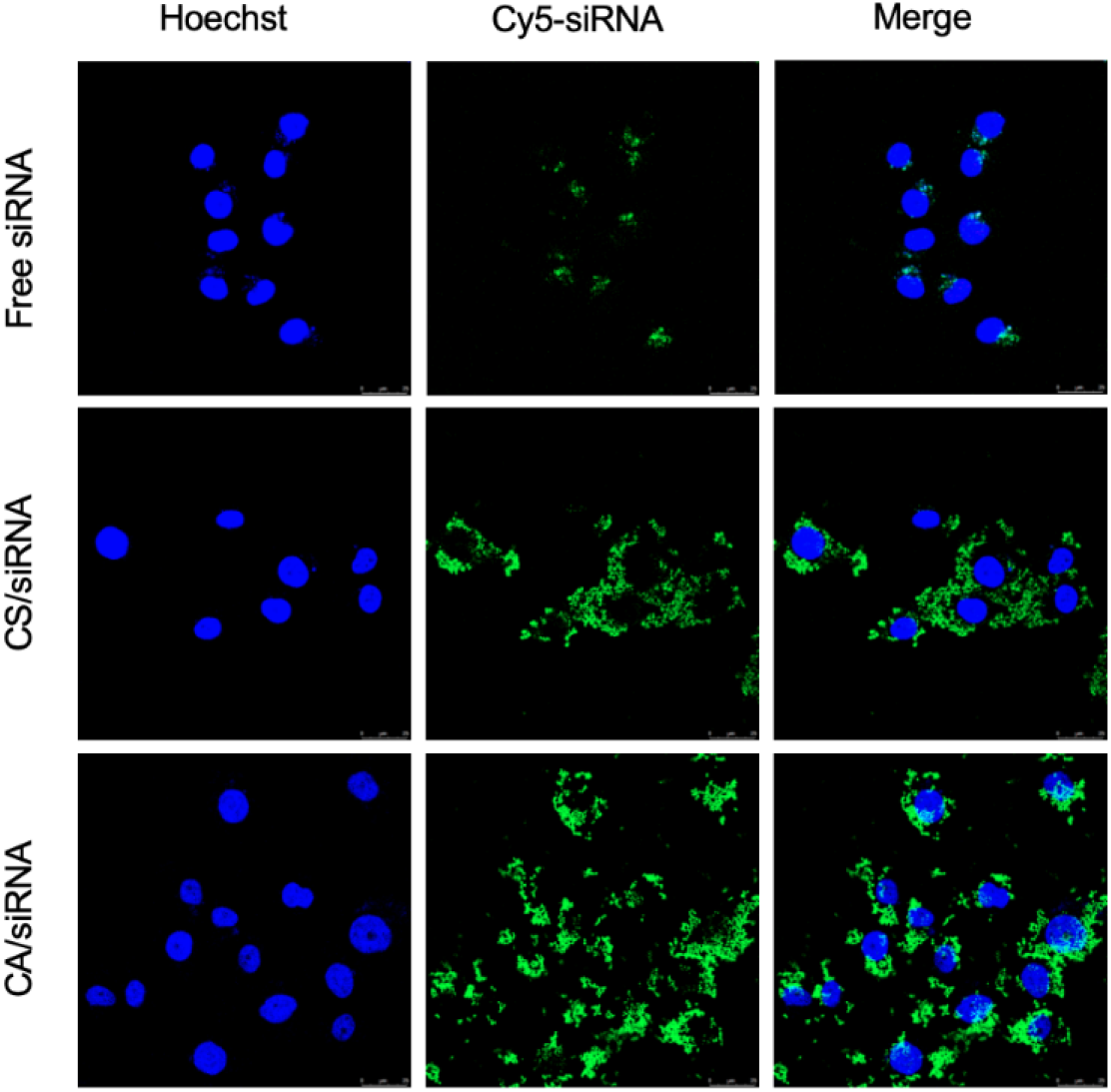
Confocal images of free siRNA, CA nanoparticles with a CA-to-siRNA weight ratio of 10, and Cs nanoparticles with a Cs-to-siRNA weight ratio of 10.

The better ability to enter cells for CA nanoparticles do not come from the size effects. Weight ratio 5 CA carriers are smaller than weight ratio 5 Cs carriers, and they are bigger than weight ratio 10 Cs carriers. At the same time, weight ratio 10 CA carriers are bigger in size than both weight ratio 5 Cs carriers and weight ratio 10 Cs carriers. The trend in size differences do not follow the trend in fluorescence intensity differences, and it suggests changing the nanoparticle sizes would not remarkably uplift the cellular uptake.

### 3.5 Cellular Intake with Confocal images

Confocal images can directly map out where the siRNA and nanoparticles are located, whether inside the cells, on the surface of the cells or merely in the medium. Hoechst is a dye that specifically combines inside the cells. The measurement of this channel can show where the inside of the cells is. Cy5 is a stable and bright dye that was previously synthesized onto DNA and RNA to locate them under microscope.

For both free siRNA and Cs nanoparticles, Cy5 and Hoechst barely overlap, suggesting that siRNA is not found inside cells. For CA nanoparticle, Cy5 and Hoechst overlap, suggesting that siRNA is inside each and every cell. This indicates that CA nanoparticles is good at penetrating cell membrane, and that the presence of L-arginine allows more siRNA to get into cells.

### 3.6 Knockdown efficiency

siRRM2 was used for silencing RRM2 mRNA. siNC does not match with RRM2 mRNA, and was thus used as the control group. Measurement in untreated cells is rescaled to be 100 as the reference. The relative RRM2 mRNA level of cells infected by free siRRM2 is only slightly below that of the untreated cells, suggesting that free siRRM2 is not efficient in knowcking down its targeted mRNA, as expected. The RRM2 mRNA level is considerably lower in cells infected by Cs-siRRM2 complexes than in cells infected by Cs-siNC complexes. The difference in their knockdown efficiency suggests that both siRRM2 and siNC have entered cells, implying that the Cs carrier makes it relatively easier for siRNA to penetrate cell membrane. The RRM2 mRNA level is significantly lower in cells infected by CA-siRRM2 compared with cells infected by CA-siNC, suggesting success in transporting siRNAs into cells. Contrasting the relative RRM2 mRNA levels of cells infected by free siRRM2 and cells infected by CA-siRRM2, the mRNA knockdown by CA-siRRM2 is remarkably more successful. This implies that the CA carrier significantly increases the amount of siRRM2 inside cells and immensely facilitates the knockdown of RRM2 mRNA.

Under the visible light, the microscope images in Figure 7 shows that there are significantly fewer live cells in the CA-siRRM2 group, compared with all other groups, especially the untreated and the free siRRM2 group. This shows that the CA construct is exceptionally effective in bringing siRRM2 into effect and killing predominantly more cells

**Figure 7.**
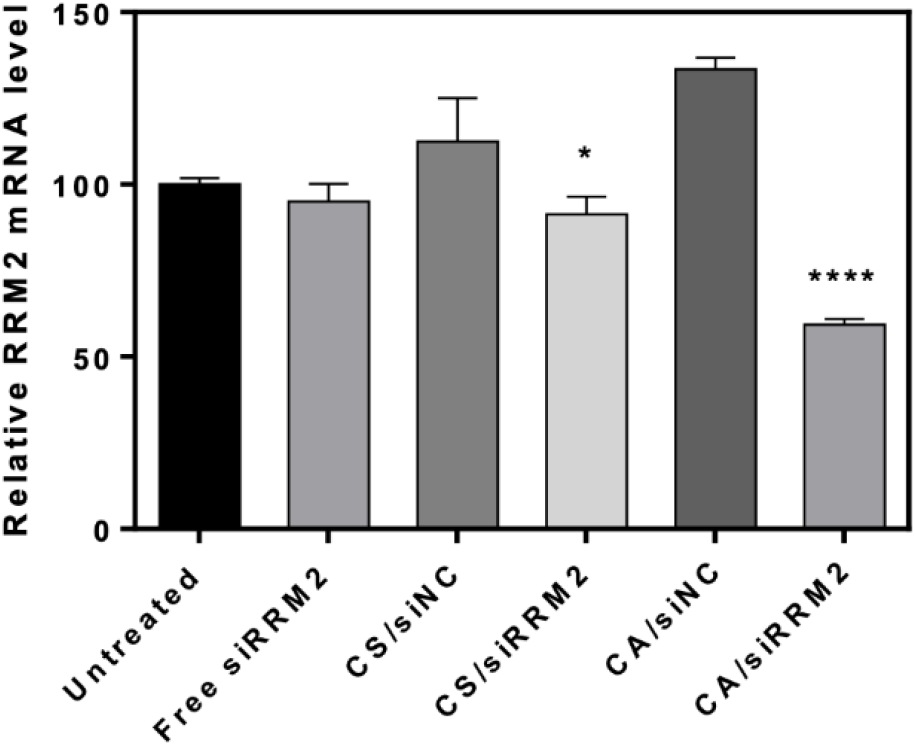
Quantification of RRM2 mRNA level after transfected by free siRRM2, Cs-siNC, Cs-siRRM2, CA-siNC, and CA-siRRM2.

**Figure 8.**
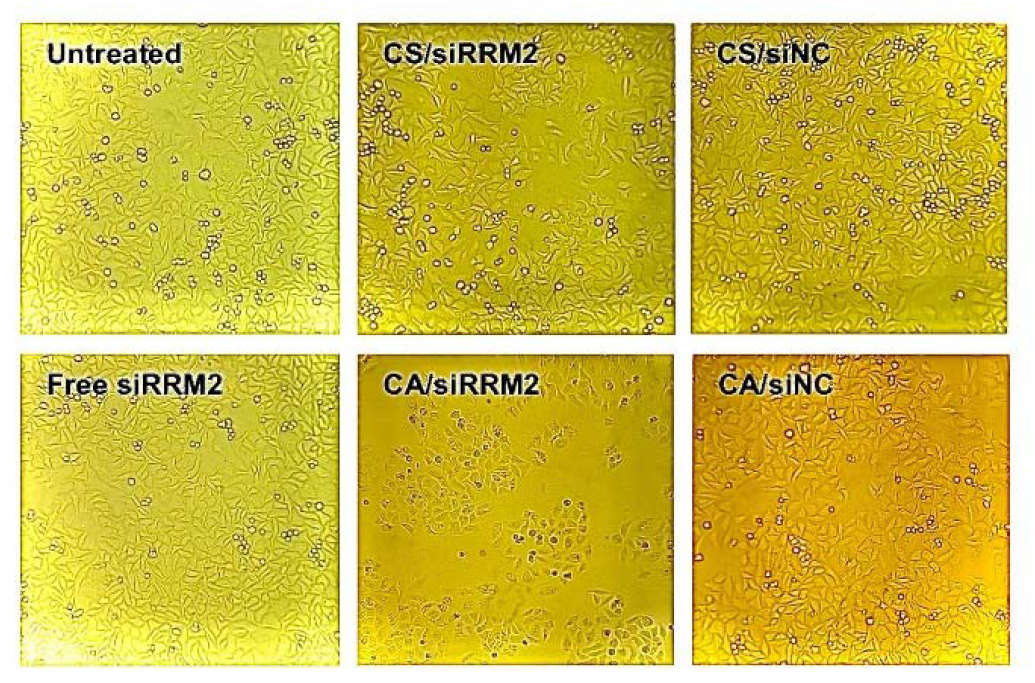
Images of H1229 after transfected by free siRRM2, Cs-siRRM2, Cs-siNC, CA-siRRM2 and CA-siNC.

## 4. Discussion

Bearing in mind the high mistargeting rate of chemotherapy, as well as the severe immune responses incurred by viral vectors, we conducted this study seeking to find a nucleic acid carrier superior to the currently used viral vector, improving upon both the efficacy and safety of existing gene therapy technologies. With its high cellular uptake efficiency, knock-down efficiency of targeted mRNA, and cytotoxicity to tumor cells, the Cs-Arg compound is a desirable vector for siRNA delivery.

This study showed RNA interfering is feasible in cancer treatment, as our siRNA competently hampered the tumor cells survival rate, the capability of our artificially synthesized Cs-Arg carrier, as it largely facilitates the uptake of siRNA in vitro, as well as the high possibility of large-scale production, as the synthesis process is extremely simple, demanding nothing more than a reaction environment with the right pH and temperature.

To further investigate the efficacy of our Cs-Arg carrier, still more tests are necessary. Previous studies have pointed out that chitosan-based delivery system tends to be limited in terms of its low water solubility, charge deduction at physiological pH and ineffective targeting capability[24]. Hence, in vivo tests are needed to examine the nanoparticle’s performance in clinical applications. There are a number of possible obstacles to which we must pay attention, such as whether the Cs-Arg compound still keeps its charge in the internal environment of organisms, where pH often strays from the optimal weak-acid range demanded by Cs, whether the addition of targeting ligands is needed to improve the specificity of the drug, or whether surface decoration would make Cs toxic, or whether there is still space of improvement in terms of how well the nucleic acid is released after endosomal uptake. 1

## 5. Conclusion

siRNA carrier was synthesized by attaching L-arginine to chitosan, promising a high cellular uptake efficiency. The results indicated success. Presence of the Chitosan-Arginine carrier immensely boosted the internalization success of siRNA. Besides, Cs-Arg-siRNA nanoparticles effectively inhibits tumor cell growth, knocking down 70% of the targeted mRNA and significantly reducing the number of live cells 4 hours after drug use.

## Footnotes

1 https://www.ncbi.nlm.nih.gov/pmc/articles/PMC6627531/

## Notes

### Competing Interest Statement

The authors have declared no competing interest.

